# Incorporating outlier information into diffusion MR tractogram filtering for robust structural brain connectivity and microstructural analyses

**DOI:** 10.1101/2021.06.09.447697

**Authors:** Viljami Sairanen, Mario Ocampo-Pineda, Cristina Granziera, Simona Schiavi, Alessandro Daducci

## Abstract

The white matter structures of the human brain can be represented using diffusion-weighted MRI tractography. Unfortunately, tractography is prone to find false-positive streamlines causing a severe decline in its specificity and limiting its feasibility in accurate structural brain connectivity analyses. Filtering algorithms have been proposed to reduce the number of invalid streamlines but the currently available filtering algorithms are not suitable to process data that contains motion artefacts which are typical in clinical research. We augmented the Convex Optimization Modelling for Microstructure Informed Tractography (COMMIT) filtering algorithm to adjust for these signal drop-out motion artifacts. We demonstrate with comprehensive Monte-Carlo whole brain simulations and in vivo infant data that our robust algorithm is capable in properly filtering tractography reconstructions despite these artefacts. We evaluated the results using parametric and non-parametric statistics and our results demonstrate that if not accounted for, motion artefacts can have severe adverse effect in the human brain structural connectivity analyses as well as in microstructural property mappings. In conclusion, the usage of robust filtering methods to mitigate motion related errors in tractogram filtering is highly beneficial especially in clinical studies with uncooperative patient groups such as infants. With our presented robust augmentation and open-source implementation, robust tractogram filtering is readily available.

**Highlights:**

- We present a novel augmentation to tractogram filtering method that accounts for subject motion related signal dropout artefacts in diffusion weighted images.
- Our method is validated with realistic Monte-Carlo whole brain simulations and evaluated with in vivo infant data.
- We show that even if data has 10% of motion corrupted slices our method is capable to mitigate their effect in structural brain connectivity analyses and microstructural mapping.

## 1. Introduction

Diffusion-weighted magnetic resonance imaging (dMRI) of the human brain (Basser et al., 1994) has various applications ranging from early clinical stroke diagnostics (Horsfield and Jones, 2002) to investigations of the microstructural properties of the tissue (Alexander et al., 2019; Novikov et al., 2019) and structural brain connectivity mapping (Griffa et al., 2013; Delettre et al., 2019; Zhang et al., 2021). The latter two are gaining popularity in clinical research (Kamiya et al., 2020) to investigate various brain diseases and neurological conditions of adults (Fieremans et al., 2013; Benitez et al., 2014) and development of the growing brain in children and adolescents (Genc et al., 2017; Huber et al., 2019). Furthermore, with the latest advances in automatic brain segmentation with tools like Infant FreeSurfer (Zöllei et al., 2020), it is likely that the amount of brain connectivity studies of infants (Kunz et al., 2014; Pannek et al., 2018; Pecheva et al., 2019) will grow in the near future too.

The clinical dMRI research comes with its own challenges to solve, with one most difficult being the patient motion. The subject motion can be unavoidable when imaging infants or patients in discomfort or pain, resulting in complex missing data problems (Sairanen et al., 2017, 2018). In short, rapid subject motion can result in slicewise signal dropout artefacts. For readers interested in why this happens, we recommend the section “Origin of the dropout” by (Andersson et al., 2016). Therefore, the processing of the motion corrupted images requires specialized algorithms and robust methods to minimize motion induced bias in the results. While robust modeling has been considered in the contexts of diffusion and kurtosis tensor estimations (Chang et al., 2005, 2012; Tax et al., 2015) as well as in higher order models (Pannek et al., 2012) that could be used for tractography purposes, it has not been investigated thoroughly in the context of the brain structural connectivity analyses.

Structural brain connectivity analyses are based on the rapidly developing dMRI tractography (Basser et al., 2000) algorithms that represent the brain white matter structures with streamlines. These streamlines can be used to investigate which gray matter regions might have a structural link. In general, the tractography algorithms are sensitive but they lack specificity and they find great number of false streamlines connections (Thomas et al., 2014; Maier-Hein et al., 2017). This means that two gray matter regions could be linked by tractography streamlines despite that the brain tissue does not form a true structural link. This is a known issue in structural connectivity analyses (Drakesmith et al., 2015; Zalesky et al., 2016; Yeh et al., 2020) to which tractogram filtering has been proposed as one solution. Tractogam filtering can be achieved with different approaches (Zhang et al., 2021), one being the Convex Optimization Modelling for Microstructure Informed Tractography (COMMIT) (Daducci et al., 2015) which we will use in this study to demonstrate possible effects of subject motion to the filtering and microstructural mapping as well as how it can be accounted and corrected for.

There are three alternative post-scan approaches to address the outliers caused by the subject motion. The first approach is to find outliers in dMRI data manually or automatically with statistical methods or deep-learning and simply exclude the artefactual dMRI data or even the whole subject from the analysis (Oguz et al., 2014; Samani et al., 2019). The second approach is to use a model to predict what the measurements should look like, locate the outliers based on differences to model predictions and replace them with these predictions if differences are deemed large enough (Lauzon et al., 2013; Andersson et al., 2016). The third approach is to detect the outliers, but instead of replacing or completely excluding them, their weight is reduced in all subsequent model estimation steps (Sairanen et al., 2018).

Manual outlier detection can be laborious and excluding whole subjects from clinical studies with relatively small number of participants might not be the optimal choice. The outlier replacement approach relies on the quality and robustness of the chosen model and method to represent the measured dMRI signal. If multiple dMRI measurements are corrupted by motion artifacts, this initial modeling and prediction step can fail altogether (Sairanen et al., 2018). Even in the best case, the replaced data points are simply interpolations based on the chosen model and the data points used in the modeling therefore it cannot increase the available information but leads to increased error propagation due to subsequent model fittings. The third approach, on the contrary, enables quantifying the amount of the motion corrupted data and versatile subsequent modeling and analysis options therefore being optimal for our purposes. In Discussion section “Robust modeling or outlier replacement” we provide further reasoning why we promote the use of robust methods over replacement in dMRI.

While weighted and robust modeling has been implemented before, they have mostly been used outside the scope of tractogram filtering. For example, in diffusion tensor modeling weighted linear least squares is typically the fastest and most robust estimator (Veraart et al., 2013; Tax et al., 2015; Sairanen et al., 2018). Robust modeling has been proposed for higher order models as well (Pannek et al., 2012). In the context of tractogram filtering, weighted cost functions have been introduced earlier in e.g. SIFT (Smith et al., 2013, 2015), but it has only been evaluated with voxels affected by partial voluming. SIFT algorithm states that their ’processing mask’ is ’the square of the estimated white matter partial volume fraction’ - which indeed should be beneficial in the case of partial voluming. However, the approach in SIFT does not account for outliers that are randomly occurring in the measurements as our newly proposed augmentation to COMMIT does.

In this work, we propose a robust augmentation to the COMMIT filtering algorithm (Daducci et al., 2015) that accounts for the unreliability of the original measurements. V Sairanen et al.: *Preprint submitted to Elsevier* Page 2 of 16 We detail the theoretical changes to the algorithm as well as provide open-source code^1^ of its implementation. We refer to this new method as COMMIT_r throughout this manuscript. To evaluate the method, we use the data from the Human Connectome Project (HCP) (Van Essen et al., 2013) as a base for thorough Monte-Carlo simulations which emulate various motion induced artifacts in synthetic but realistic whole brain data. Synthetic data provides the necessary baseline that can be used to isolate the bias arising from subject motion from noise effects in structural connectivity analyses as well as how well motion artefacts can be amended using our robust augmentation. In the context of this study, the measurement unreliability is associated with outliers due to subject motion. However, it can readily be utilized to correct for measurements that are affected by partial voluming, as our preliminary results have demonstrated earlier (Sairanen et al., 2021).

## 2. Materials and Methods

### 2.1. Implementation

We augmented the original cost function of COMMIT (Daducci et al., 2015) with a voxelwise weighting factor **W** that we used to down weight measurements that have decreased reliability due to subject motion or any other reason. The original COMMIT is based on a minimization of the difference between the original measurements and a forward model prediction. The forward model prediction is calculated by fitting a chosen microstructural model for each streamline in every voxel. COMMIT assigns a weight to each streamline that tells how much that streamline contributes to the predicted signal. These streamline contribution weights are iteratively updated until the difference between the measurements and this prediction converges to a minimum. Any streamline with contribution of zero is then removed as an implausible streamline (i.e. not compatible with the measured signal).

If part of the measurements are artefactual due to subject motion or any other reason, the original COMMIT algorithm could converge to an incorrect solution. To avoid this and decrease the impact of these artefactual measurements, we propose the robust cost function shown in eq. 1. The weighting factor (**W**) is used to multiply the difference between the original measurements (**y**) and the product of model design matrix (**A**) and estimated model coefficients (**x**) in the minimization problem. Our proposed idea is further illustrated in Fig. 1 with a simple toy example. In future, these reliability weights **W** could be iteratively updated along with the model coefficients **x** to help in estimating model coefficients in voxels that do not fit to the chosen model perfectly due to heart beat related pulsation or other uncertainties.

**Figure 1:**
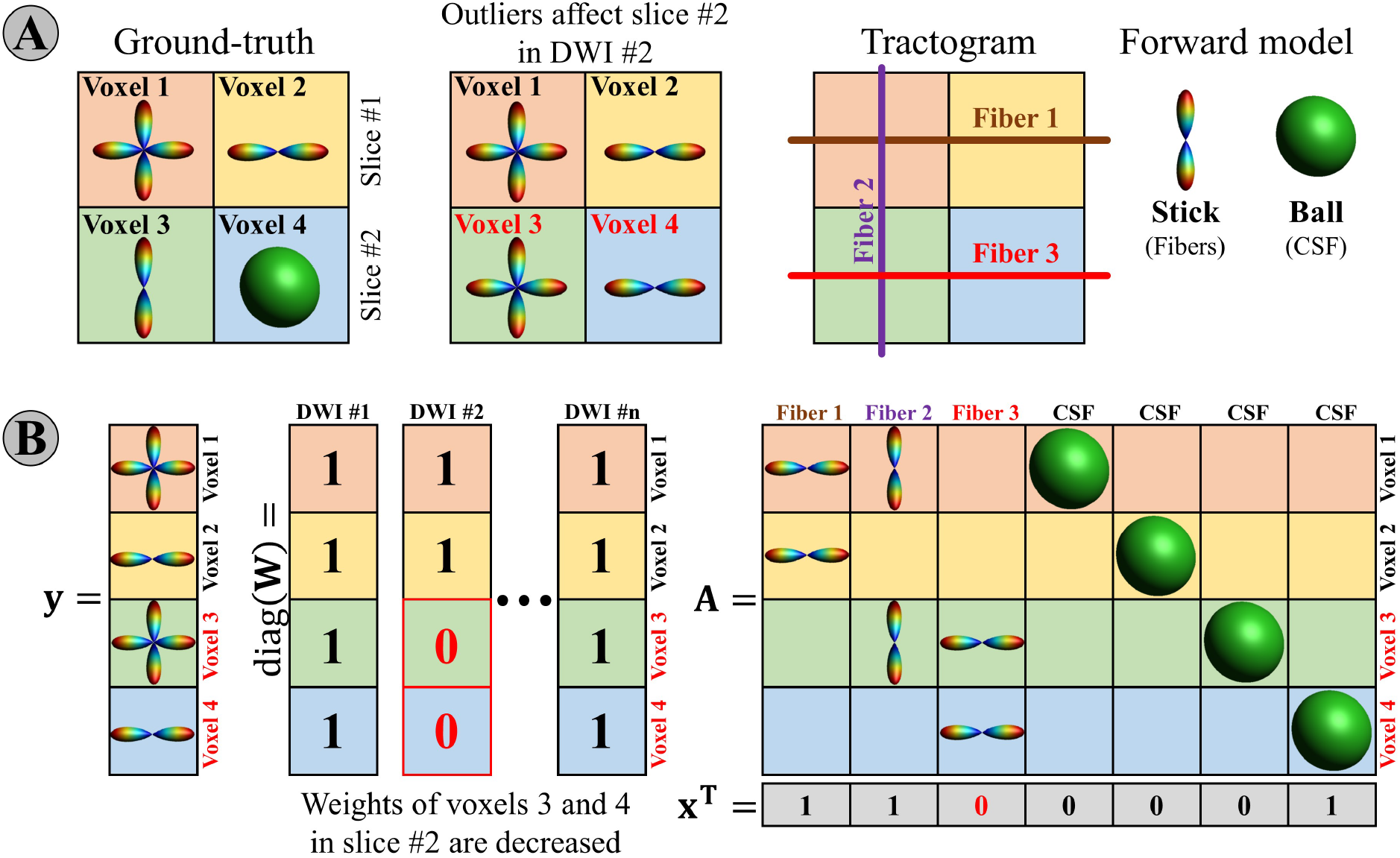
A toy example to illustrate the augmentation of COMMIT framework with the measurement reliability based weighing. (**A**) A synthetic phantom consists of two slices both consisting of two voxels. For visualization purposes the 4th dimension i.e. diffusion weighted signals is omitted and observed signal is visualized using a fiber orientation distribution (FOD). To illustrate the subject motion artefact, the second slice in DWI #2 is affected by slicewise artefact and FOD of corresponding voxels 3 and 4 is biased. In tractography, this is seen as an implausible streamline Fiber 3. (**B**) Data is vectorized to visualize the linear model problem. Vector **y** contains the simulated signals, diagonal of matrix **W** contains the reliability weights that are decreased for voxels 3 and 4 in DWI #2, **A** is the design matrix that depicts the modeled compartments (e.g., stick for streamlines and ball for isotropic compartments), and **x** contains the streamline wise contributions. With the successful downweighing, the contribution of Fiber 3 is set to zero and the implausible streamline is thus removed.

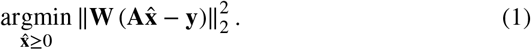

The robust cost function in eq. 1 is intended to be used with outlier detection with tools such as SOLID (Sairanen et al., 2018). SOLID detects slicewise outliers based on robust statistical analysis of the original dMRI data and can be used either to exclude outliers or down weight them depending how strong outliers are. This down weighting scheme is likely a better option to outlier replacement that is proposed in earlier studies (Lauzon et al., 2013; Andersson et al., 2016). If the outlier is replaced with a prediction from a tensor or a gaussian model, then COMMIT would try to minimize the difference from those model predictions to its own model prediction. Since these models can be different and therefore capture different details of the dMRI signal, it is more straightforward to use robust modeling with the proposed weighted cost function. For interested readers, we provide more reasoning for this claim in the Discussion section “Robust modeling or outlier replacement”.

### 2.2. Simulations

To investigate the outlier effect on the tractogram filtering, we developed a comprehensive Monte-Carlo simulation pipeline delineated in Fig. 2. Simulations were based on T1-weighted and dMRI data from the HCP subject 100308 which were processed with current state-of-the-art methods (Van Esen et al., 2013). We do not expect or imply that this ground truth connectivity matrix depicted in Fig. 3 would represent the true structural connections in a human brain. It simply provides us the necessary ground truth connectivity that we can use to evaluate the noise and outlier effects in the Monte-Carlo simulations with more realistic picture of the whole brain than typical fiber phantoms as it contains realistic brain structures such as kissing an crossing fibers as well as the modeled partial voluming effects.

**Figure 2:**
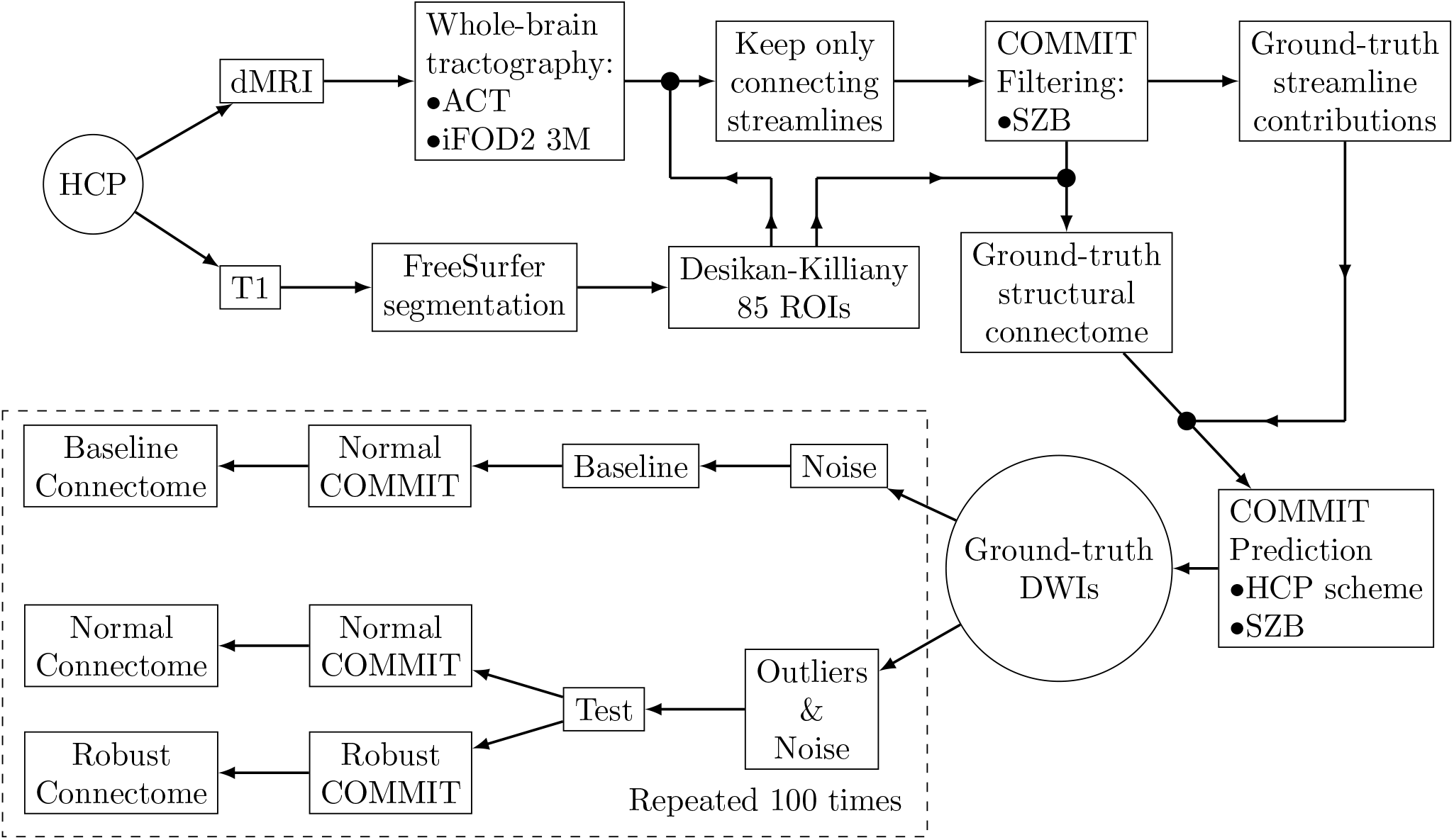
A flow chart describing how the whole-brain simulations were obtained from the HCP dataset. The dMRI and T1-weighted data were used to obtain the ground truth connectome from which the ground truth dMRI signals were predicted using normal COMMIT forward modeling. The ground truth data was used to perform 100 Monte-Carlo simulations to evaluate the effects of noise and outliers to the structural brain connectome. The Monte-Carlo iteration setups shown inside the dashed rectangle were repeated for outlier percentages 5% and 10% both with the uniform and clustered outlier schemes.

**Figure 3:**
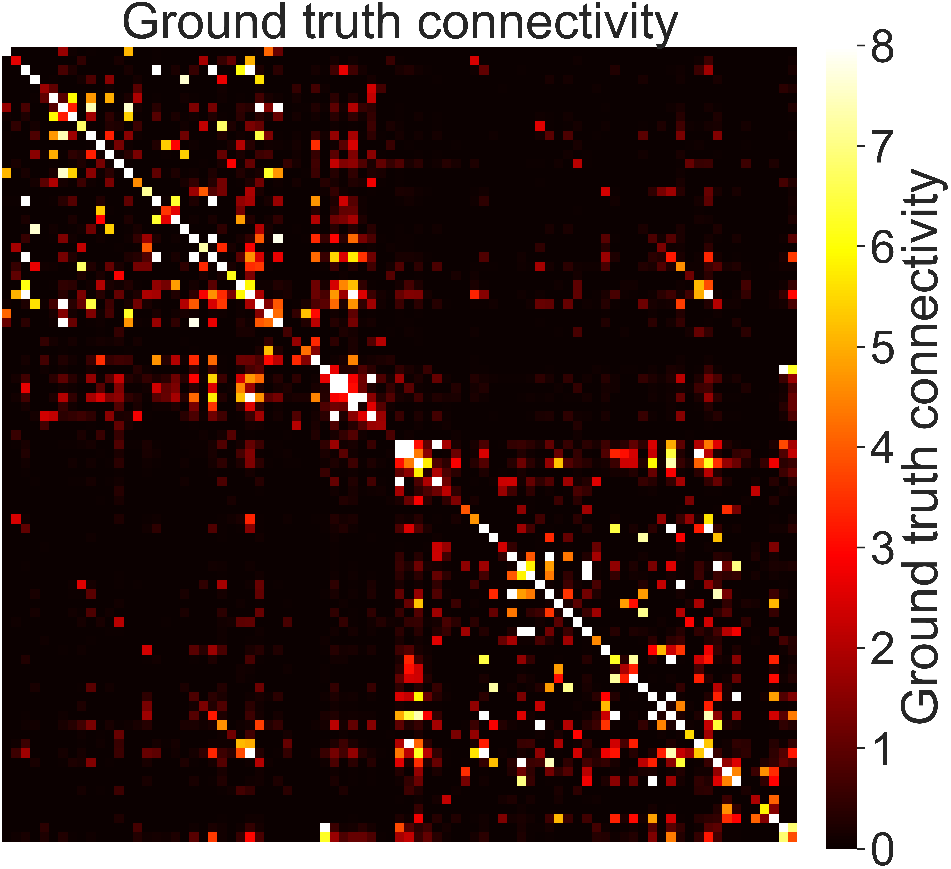
The ground truth connectivity matrix used in this study was based on one subject. While, this connectivity matrix might not represent the real human brain connections, it provided the necessary ground truth control for our Monte-Carlo simulations. Deviations from this network in the simulations would be due to noise, outliers, or both.

#### Ground truth data

We segmented the T1-weighted HCP data with FreeSurfer (Fischl, 2012) to obtain 85 regions-of-interests (ROIs) based on the Desikan-Killiany atlas (Desikan et al., 2006). Instead of the full brainstem, we used only its inferior part of medulla as the last ROI. We used these brain segments to compute the ground truth connectivity matrix as well as to ensure that we used only the connecting streamlines in our analyses.

To calculate a whole brain tractogram from the HCP dMRI data, we used the anatomically constrained probabilistic tractography (iFOD2) (Tournier et al., 2010; Smith et al., 2012) implemented in MRTrix3 software (Tournier et al., 2019). We used the white matter mask as a seed region for three million streamlines. The tracking parameters were: step size 0.5, turning angle 45 degrees, min length value 5, max length 250, cutoff value 0.05, trials number 1000. Finally, we removed all non-connecting streamlines based on the 85 ROI segmentation of T1-image.

For ground truth tractogram filtering, we used the original COMMIT (Daducci et al., 2015) because data did not contain slicewise outliers. We chose the stick-zeppelin-ball (SZB) as the forward model with following parameters: 1.7 · 10^−3^*mm*^2^/*s* for parallel stick and zeppelin diffusivities, 0.61·10^−3^*mm*^2^/*s* for perpendicular zeppelin diffusivity, and two ball compartments of 1.7 · 10^−3^*mm*^2^/*s* and 3.0 · 10^−3^*mm*^2^/*s* to account for partial volumeing with gray matter as well as in cerebrospinal fluid (Panagiotaki et al., 2012).

The filtered tractogram was used to form the ground truth connectivity matrix with the information from T1-segmentation (Fig. 3). The network edges in the ground truth connectivity matrix were defined as the sum of the COMMIT streamline weights multiplied by the length of the tract and normalized by the average tract length between each gray matter parcellation as was done in (Schiavi et al., 2020). We combined this information with the final streamline contributions to form the synthetic whole brain prediction of dMRI data using the HCP’s three-shell gradient scheme. This produced 270 noise free diffusion-weighted whole brain images that we used as a ground truth for our Monte-Carlo simulations.

#### Monte-Carlo data

Our Monte-Carlo simulations were based on the ground truth synthetic whole brain dMRI data obtained from HCP subject. We split the simulations in two groups: Baseline and Test. Baseline group provides the means to evaluate the pure noise effects on the connectome whereas Test group provides the means to isolate and evaluate the outlier effects. Network analyzes used for Test group were identical with the ground truth case.

In Baseline group, random Rician noise was added before repeating the normal COMMIT filtering with the original non-filtered but connecting streamlines. The Rician noise had signal-to-noise ratio of 20 based on the non-diffusion weighted signal which is roughly similar with signal-to-noise ratios in clinical research. We used the same filtering parameters that were used to form the ground truth data. This process was repeated to obtain 100 whole brain baseline images and connectomes.

In Test group, outliers were introduced to the data before adding the same Rician noise that was used for the Baseline group. Test group was filtered with both the normal COMMIT as well as the proposed robust COMMIT_r using the same streamlines and parameters that were used for the Baseline group. This process was repeated to obtain 100 whole brain test images with outliers and corresponding connectomes from normal and robust filtering methods.

The outlier selection for the Test group was done with two different schemes by replacing axial slices with signal decrease outliers in an interleaved manner to 5% and 10% of the dMRI data per shell. The first scheme represented the worst possible situation where outliers were clustered in the q-space (e.g. Fig. 4) whereas the second scheme represented the best possible situation where outliers were uniformly placed in the q-space based on their electrostatic repulsion (Sairanen et al., 2017). Futher details why we did not use purely random selection of outliers is provided in Discussion section “Robust modeling or outlier replacement”.

**Figure 4:**
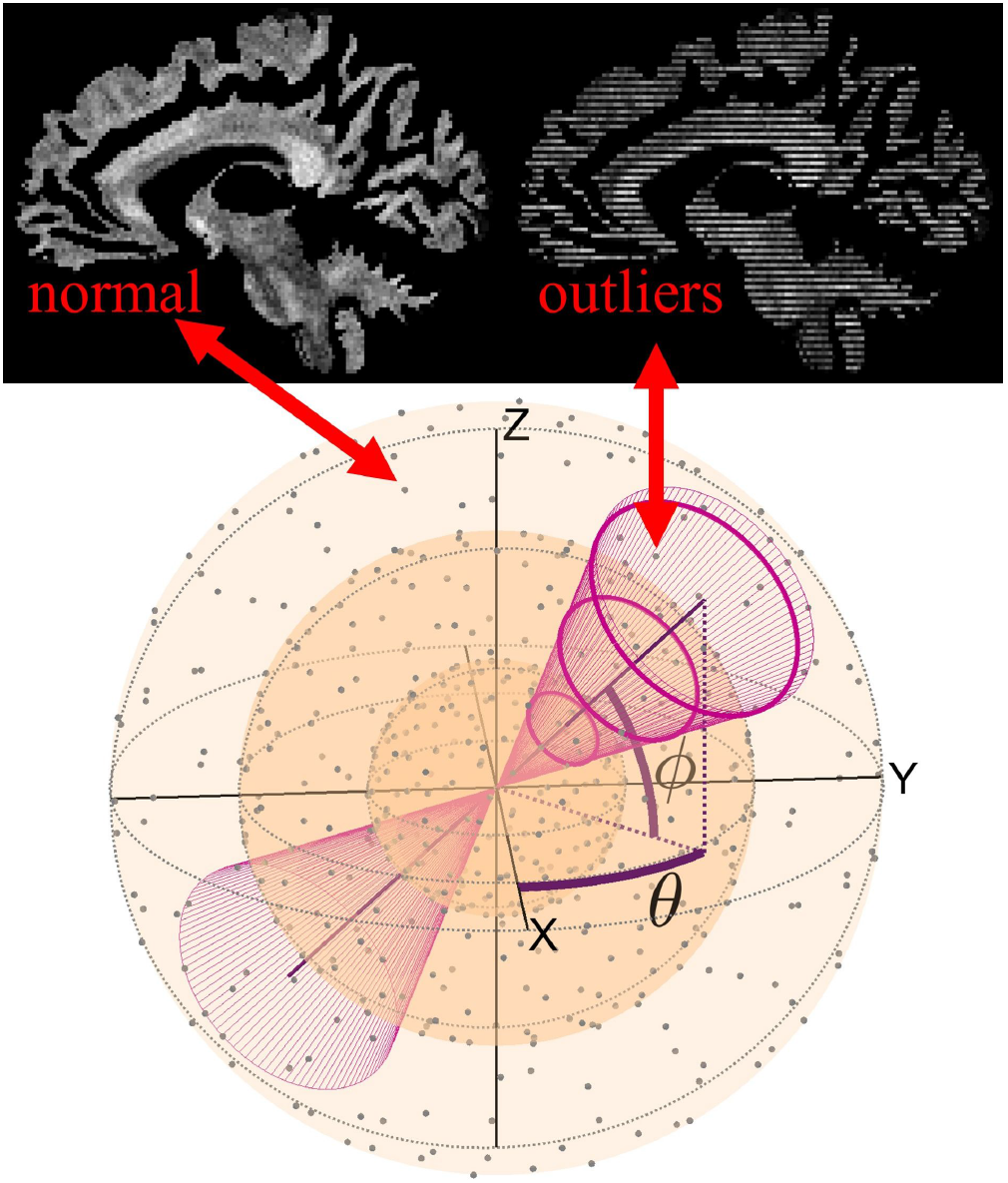
An illustration how clustered outliers were selected from all the gradient directions. The three shells used in the HCP gradient scheme are shown with the transparent spheres and gradient directions with black dots. The initial direction (*θ, ϕ*) of was selected randomly after which the opening angle of the cone was increased until wanted number of outliers from each shell remained inside it. This approach ensured the maximal gap in the *q*-space sampling and the chance to find error prone schemes.

#### Statistical analysis

We investigated global brain connectivity as well as individual network edges using analysis of variance (ANOVA) accompanied by Tukey’s honestly significant difference (HSD) test and non-parametric Friedman’s test accompanied by two-sample Kolmogorov-Smirnov tests. The reason for having these different test statistics is that outliers can lead to skewed and long tailed distributions that might not be correctly investigated solely by parametric tests. It should be noted that the added Rician noise has a non-zero positive mean value. This means that all groups are likely shifted to some direction from the ground truth prediction. This is the reason, why the Baseline group is needed as that is affected only by noise and can be used to isolate corresponding shifting effect.

While we report p-values from these tests, we argue that the effect sizes are more interesting as they describe how different the tested groups are. The effect sizes are measured using Cohen’s D for parametric tests and Kolmogorov-Smirnov statistic for non-parametric tests. The test statistics we employ are widely used and they provide information about average differences and differences in the shapes of the Monte-Carlo simulated distributions. For details about these tests, we recommend any textbook that covers parametric and non-parametric statistics such as Sheskin’s handbook (Sheskin, 2004). Having these two different tests seemed necessary during designing this study. Of course, as we did multiple tests, we also performed multiple comparison correction to all the p-values in pair-wise tests. In total, there was 80 different tests and we corrected for them using Benjamini-Hochberg False Discovery Rate (FDR) test (Benjamini and Hochberg, 1995) with alpha 0.05. However, as seen from the results later, neither mean based or distribution shape based analysis might not be sufficient to provide absolute conclusions.

### 2.3. In vivo measurements

#### Infant data

We obtained preliminary data from an on-going infant study to evaluate our method with in vivo measurements. T1-weighted image and dMRI data were obtained with 3T MRI Siemens Skyra system (Erlangen, Germany) with a 32 channel head coil. The dMRI acquisition consisted of 13 non-diffusion weighted images that were interspersed between 60 diffusion-weighted images with b-value of 750*s*/*mm*^2^ and 74 diffusion-weighted images with b-value of 1800*s*/*mm*^2^ each with uniquely oriented gradients. Bipolar gradient scheme was used to minimize geometrical distortions due to eddy currents. The image resolution was isotropic 2*mm* with 80 × 80 × 44 imaging matrix. The in-plane acceleration factor was 2 (SENSE) and multi-band acceleration factor was 2. Only anterior-posterior phase encoded images were acquired as the reverse phase encoding required manual adjustment during the scan which was deemed infeasible at the corresponding clinical scan environment. The use of infant data in this work was approved by the relevant Ethics Committee of the Helsinki University Hospital.

#### Infant analyses

We used ExploreDTI (Leemans et al., 2009) with SOLID-plugin (Sairanen et al., 2018) to simultaneously detect slice-wise outliers and to correct for subject motion and eddy currents as well as registered the data to anatomical T1-image to correct for geometrical distortions. Additionally, we used Gibbs ringing correction (Perrone et al., 2015). We did not correct for signal drift (Vos et al.) as it was not observed in the measurements.

Processing of this data was limited to specific computers in the hospital network which prevented memory demanding tasks such as segmentation with Infant Freesurfer (Zöllei et al., 2020). Problematically, the T1-image contrast of this subject was not suitable for white and grey matter segmentations using traditional options. This, unfortunately, prevented us from performing full network analyses on the infant dataset as there was no reliable way to perform gray matter segmentation and we had to content to a simpler analysis that consisted of comparing signal fraction maps between normal and robust filtering method. We obtained a WM mask from multi-shell multi-tissue constrained spherical deconvolution (Jeurissen et al., 2014) implemented in MR-Trix3 (Tournier et al., 2019) and used that as a seed mask for probabilistic whole-brain tractography (iFOD2) (Tournier et al., 2010) to generate three million streamlines.

We filtered the generated streamlines with normal COMMIT (Daducci et al., 2015) and the proposed robust COMMIT_r to evaluate the improvements in the overall fit from root mean squared error (RMSE) maps as well as to see the impact of outliers in intracellular and isotropic signal fractions. We used the stick-ball model for both filtering methods with the following parameters: 1.7 · 10^−3^*mm*^2^/*s* for parallel signal diffusivity, and 1.7·10^−3^*mm*^2^/*s* and 3.0·10^−3^*mm*^2^/*s* for the isotropic signal diffusivities.

## 3. Results

### 3.1. Simulations

We investigated the effects of noise to the structural brain connectivity by comparing Baseline group. Test groups (COMMIT and COMMIT_r) could not be directly compared to the ground truth due to Rician noise bias. With the Rician noise bias we imply the effect that adding noise with non-zero mean (Gudbjartsson and Patz, 1995) to data leads to a shift in overall baseline. Therefore, outlier effects were investigated by comparing Test groups to Baseline group. We evaluated these differences in both global connectivity matrix score as well as in network edge individually.

#### Global Connectivity

The global connectivity difference was defined as an average absolute difference between the elements upper triangle of the connectivity matrices from Monte-Carlo groups and the corresponding ground truth values. The results of this comparison calculated are shown in Fig. 5 with all violin plots being based on 100 Monte-Carlo simulations each. The noise effect on the global connectivity (Baseline) is shown with the first violin from the left, the uniform outlier effect is shown in the middle, and the clustered outlier effect is shown on the right. The percentage of outliers (5% or 10%) is shown on different sides of each violin.

**Figure 5:**
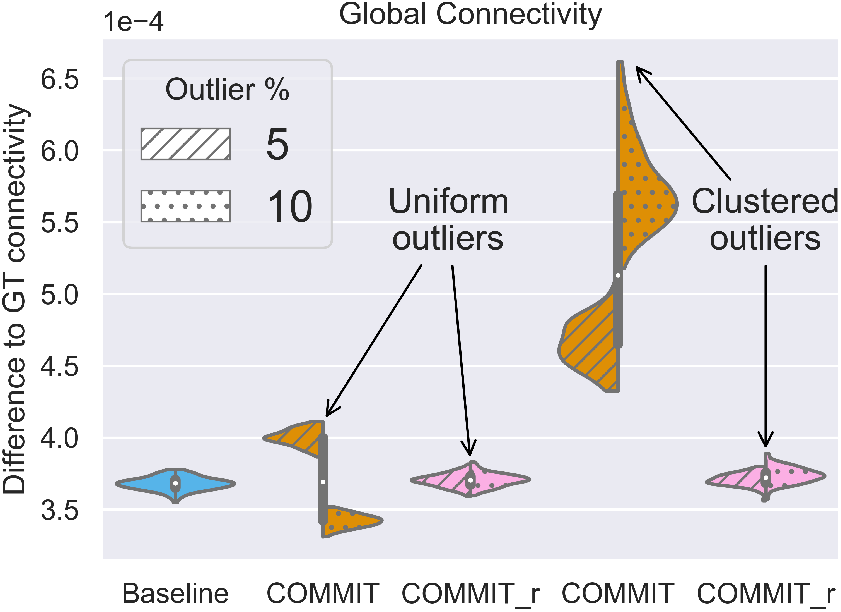
Impact of noise (Baseline) and outliers (COMMIT and COMMIT_r) to the global structural brain connectivity. Left and right sides of the violins represent simulations with 5% and 10% outliers, respectively. The y-axis indicates the distance to the ground truth as an average absolute difference. The augmented COMMIT_r shown with the pink violins produced similar distributions with the Baseline in all cases whereas the normal COMMIT shown with orange violins differs from the Baseline already in the 5% cases. As expected, the clustered outlier scheme produced the largest deviations with the highest variability in the normal COMMIT distributions. Interestingly, the uniform outlier scheme resulted two different distributions for normal COMMIT compared to the Baseline. This highlights the need for the robust processing as the exact effect of outliers can be very challenging to predict.

Both, Baseline and robust COMMIT_r produced similar global results with differences ranging from 3.5 · 10^−4^ to 4.0 · 10^−4^. This demonstrates that on average, the proposed robust filtering method is capable to mitigate the outlier effect. On the contrary, the results from normal COMMIT ranged from 3.25 · 10^−4^ to 6.5 · 10^−4^ demonstrating that outliers can have a much stronger effect than noise on the global connectivity values.

##### Parametric statistical analysis

The global connectivity differences with ANOVA detail that the group averages were statistically different with p-value less than 0.05. Tukey’s HSD test results are shown in Table 1 along with all other statistical tests results. Statisticall significant results after multiple comparison correction with FDR alpha 0.05 are shown with bolded p-values. The results depict that normal COMMIT had significantly different mean to both Baseline and COMMIT_r results and the effect sizes evaluated with Cohen’s D were systematically larger. Importantly, differences between Baseline and COMMIT_r were not statistically significant with relatively small effect sizes. These effect sizes indicate that in our realistic simulations with 5% and 10% of outliers, the average bias caused by outliers quickly increases and compromises the connectivity analyses if data is not processed robustly.

**Table 1.**
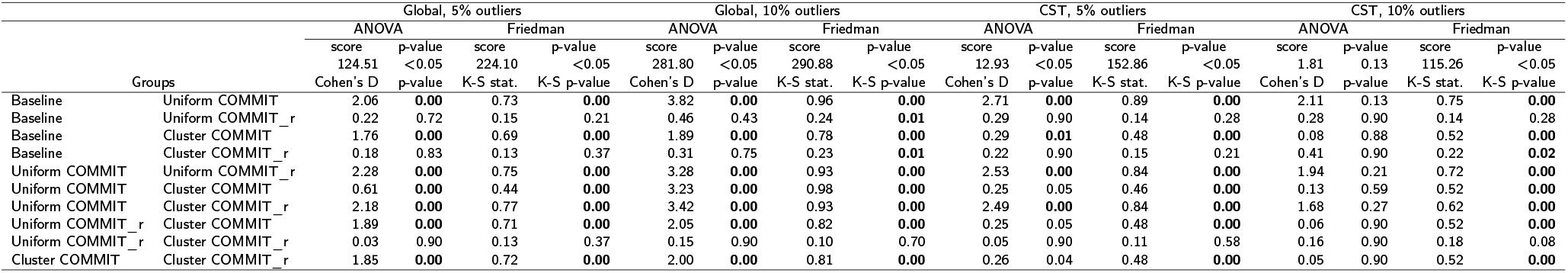
Summary of the parametric and non-parametric test results. Bolded p-values indicate statistically significant findings with FDR based correction for multiple comparisons using 0.05 alpha.

##### Non-parametric statistical analysis

The Friedman’s test reported also p-value less than 0.05 therefore providing additional support for the graphical analysis and ANOVA results. We applied the two-sample Kolmogorov-Smirnov test to detect which of the distributions were different. All these test results are reported in Table 1. Comparisons between Base line and normal COMMIT were all statistically significant with large effect sizes whereas comparisons between Baseline and COMMIT_r were not statistically significant with 5% outliers. With 10% outliers non-parametric differences between Baseline and COMMIT_r were significant but the effect size remained small.

#### Network edges

We investigated the network edge-wise differences between the Monte-Carlo connectivity matrices with parametric and non-parametric statistics as complementary information to the global results. The three violin plots in Fig. 6 depict the connectivity values from medulla to the right precentral gyrus. These streamlines are visualised in Fig. 7 and are likely a part of the corticospinal tract and therefore a known true connection. The results of the parametric and non-parametric tests performed to this network edge are depicted in Table 1.

**Figure 6:**
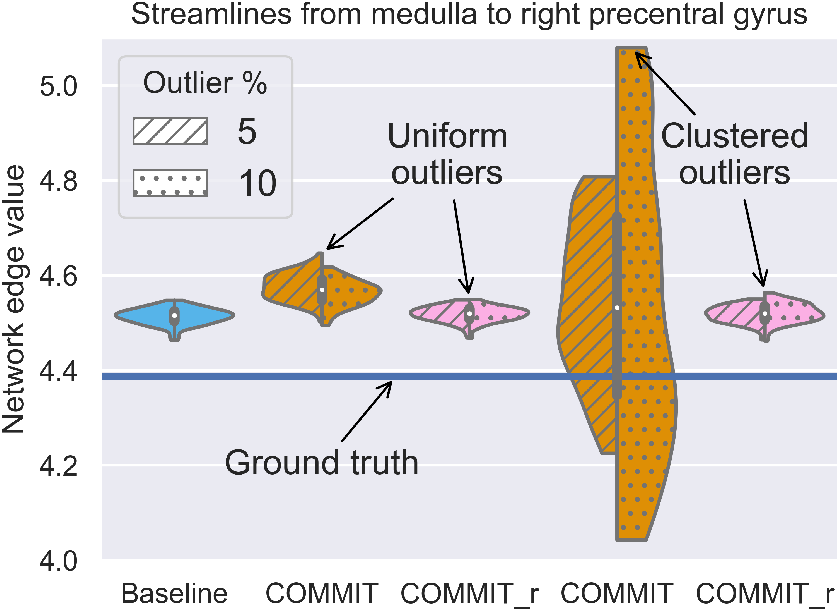
Impact of noise (Baseline) and outliers (COMMIT and COMMIT_r) to one specific network edge that represents connection between medulla and right precentral gyrus. The y-axis indicates the strength of this edge connectivity. Left and right sides of the violins represent simulations with 5% and 10% outliers, respectively. The augmented COMMIT_r shown with the pink violins produced similar distributions with the Baseline in all cases whereas the normal COMMIT shown with orange violins is heavily affected already in the 5% cases. As expected, the clustered outlier scheme produced the largest deviations with the highest variability in the normal COMMIT distributions. The normal COMMIT simulations with the clustered outlier scheme demonstrate why it is necessary to compare results against the Baseline instead of ground truth because the outlier effect can surpass the noise effect. This can lead to the shown situation where the difference to ground truth on average would be smaller due to a very wide distribution.

**Figure 7:**
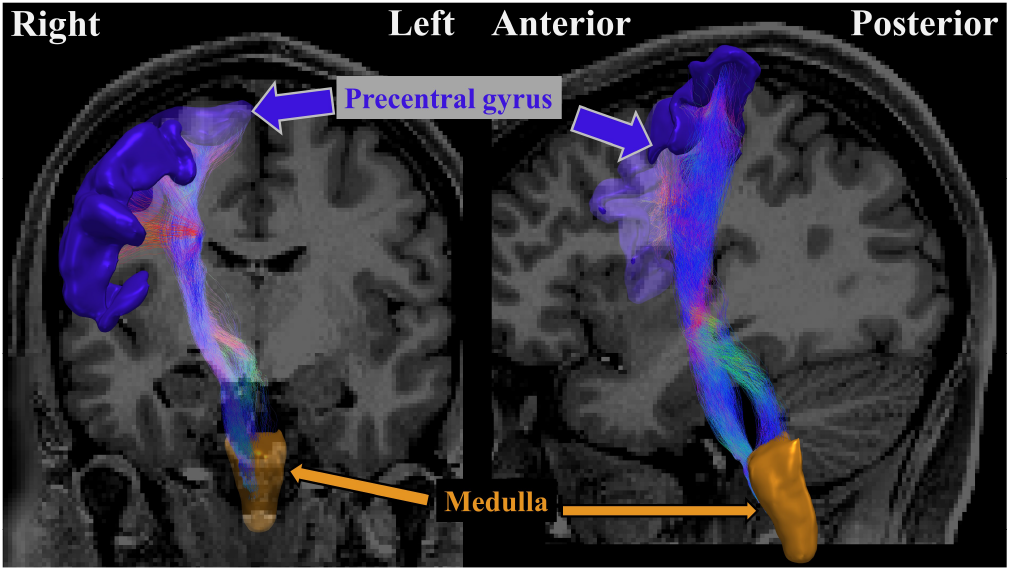
Illustration of the network edge that connects medulla to right precentral gyrus. This edge or connection was selected for closer inspection as it forms a part of the corticospinal tract which is well known true connection in a healthy human brain. Shape of the tractogram is nearly vertical therefore perpendicular to the introduced slice-wise outliers in the axial plane.

The noise effect results in a systematic over estimation of the connectivity strength as depicted by Baseline in Fig. 6. However, outliers have a more random effect depending on the affected dMRI measurements. This can either decrease or increase the connectivity strength and can counteract the noise effect. Therefore, group comparisons against Baseline were more meaningful than comparisons against the known ground truth value would be. For example, in this case the normal COMMIT produces an average connectivity strength that is closer to the ground truth than Baseline despite the distribution is wider.

##### Parametric statistical analyses

The connectivity-wise differences between Baseline and normal COMMIT as well as Baseline and robust COMMIT_r are shown in Fig. 8. The color map indicates the effect size measured with Cohen’s D. Only elements that were deemed significantly different (p-value less than 0.05) based on ANOVA and Tukey’s HSD were drawn. The comparison between Baseline and normal COMMIT resulted in more elements with significant differences than the comparison between Baseline and COMMIT_r. The effect sizes between Baseline and normal COMMIT ranged from 0 up to 3 indicating that outliers can have strong adverse effects on specific connectivity matrix elements. The overall smaller effect sizes between Baseline and robust COMMIT_r highlight that our augmentation is well capable to mitigate the outlier effects even on individual network edge level.

**Figure 8:**
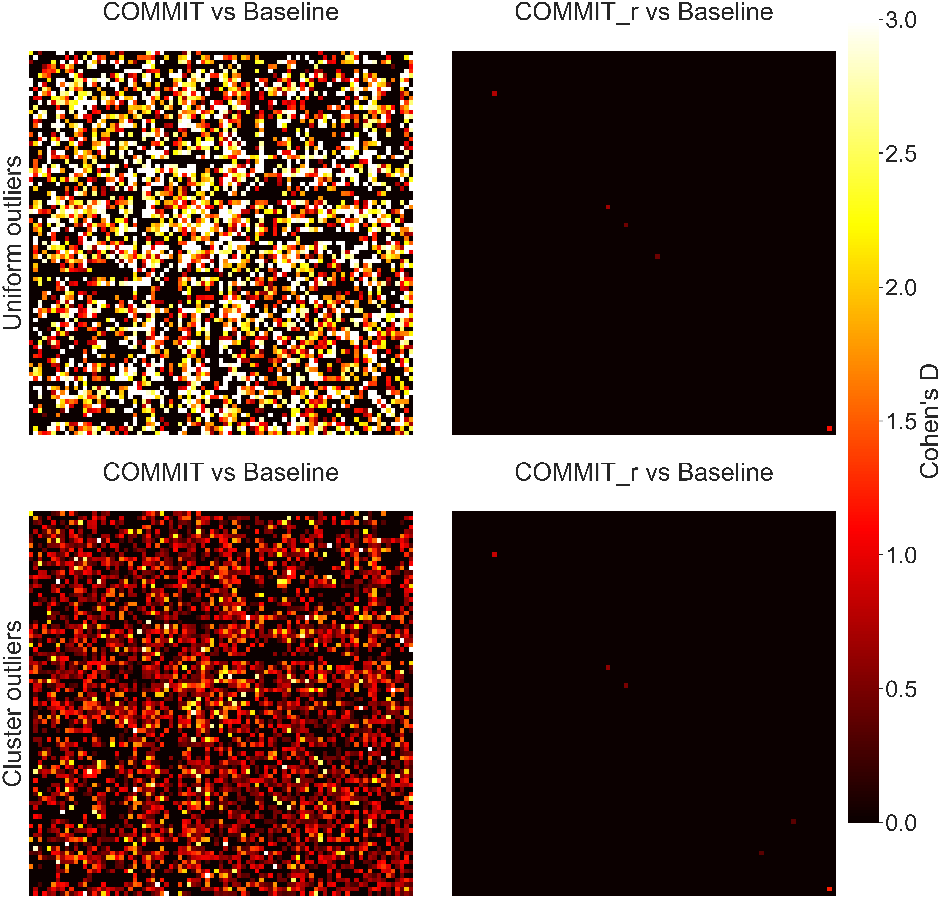
Differences in the network edges due to outliers measured using parametric statistics. The color scale indicates the average effect size calculated using Cohen’s D with unequal variances. The left and right columns show the difference from Baseline to normal and robust COMMIT. The top and bottom rows show the results from uniform and clustered outlier schemes. Robust augmentation clearly improves the COMMIT filtering if the data contains outliers as the effect sizes remain very small in all edges in both outlier schemes. While the uniform outlier scheme produced larger effect sizes for normal COMMIT than the clustered, it can be easily explained because Cohen’s D is inversely proportional to the sample variance which is very high in the clustered outlier schemes.

##### Non-parametric statistical analysis

The connectivity-wise distributional differences between Baseline and normal COMMIT as well as Baseline and robust COMMIT_r are shown in Fig. 9. The color map indicates the effect size measured with Kolmogorov-Smirnov statistic. Only elements that were deemed statistically significantly different (p-value less than 0.05) based on Kolmogorov-Smirnov tests were drawn. Similar to the parametric counterpart, the differences between Baseline and normal COMMIT were again more frequent than differences between Baseline and robust COMMIT_r. Also the effect sizes between Baseline and normal COMMIT ranged from 0 to nearly 1 which is the maximum of the used statistic. This indicates that outliers can lead to very large distributional differences. The differences between Baseline and robust COMMIT_r remained relatively small with effect sizes ranging from 0 to 0.2.

**Figure 9:**
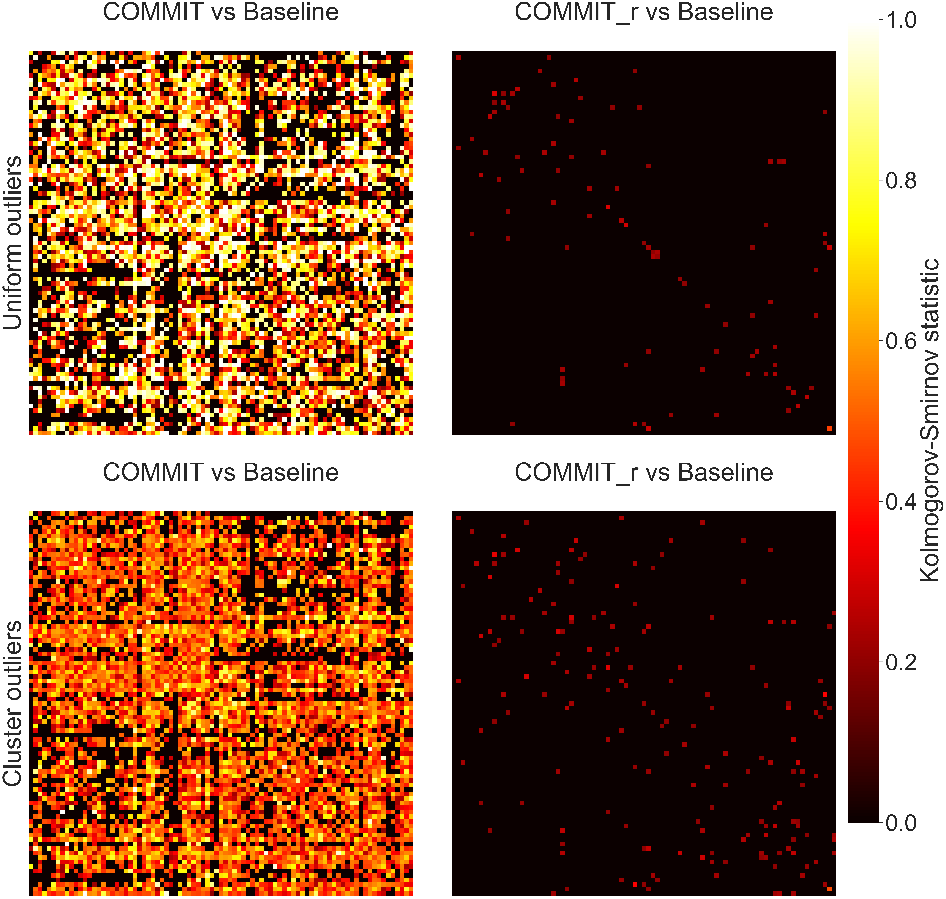
Differences in the network edges due to outliers measured using non-parametric statistics. The color scale indicates the average effect size calculated as Kolmogorow-Smirnov statistic. The left and right columns show the difference from Baseline to normal and robust COMMIT. The top and bottom rows show the results from uniform and clustered outlier schemes. Robust augmentation produces smaller effect sizes which means that the distributions between the Baseline and COMMIT_r were very similar in both outlier schemes. For normal COMMIT, the uniform outliers produced larger effect sizes than the clustered. This is the outcome of the cumulative distribution function based statistics where the uniform outliers results in distributions with a high precision but low accuracy which do not overlap with the Baseline whereas the clustered outlier results in distributions with a very low precision but moderate accuracy which do overlap with the Baseline.

### 3.2. In vivo measurements

Besides tractogram filering, we calculated the intracellular and isotropic signal fractions calculated using the COMMIT framework (Daducci et al., 2015) and the proposed robust COMMIT_r. Fig. 10 shows the results for outlier detection, RMSE, and signal fraction maps obtain from the infant data. On average, the amount of missing data i.e. how much confidence in fitting was decreased per slice position ranged from 5% to 19%.

**Figure 10:**
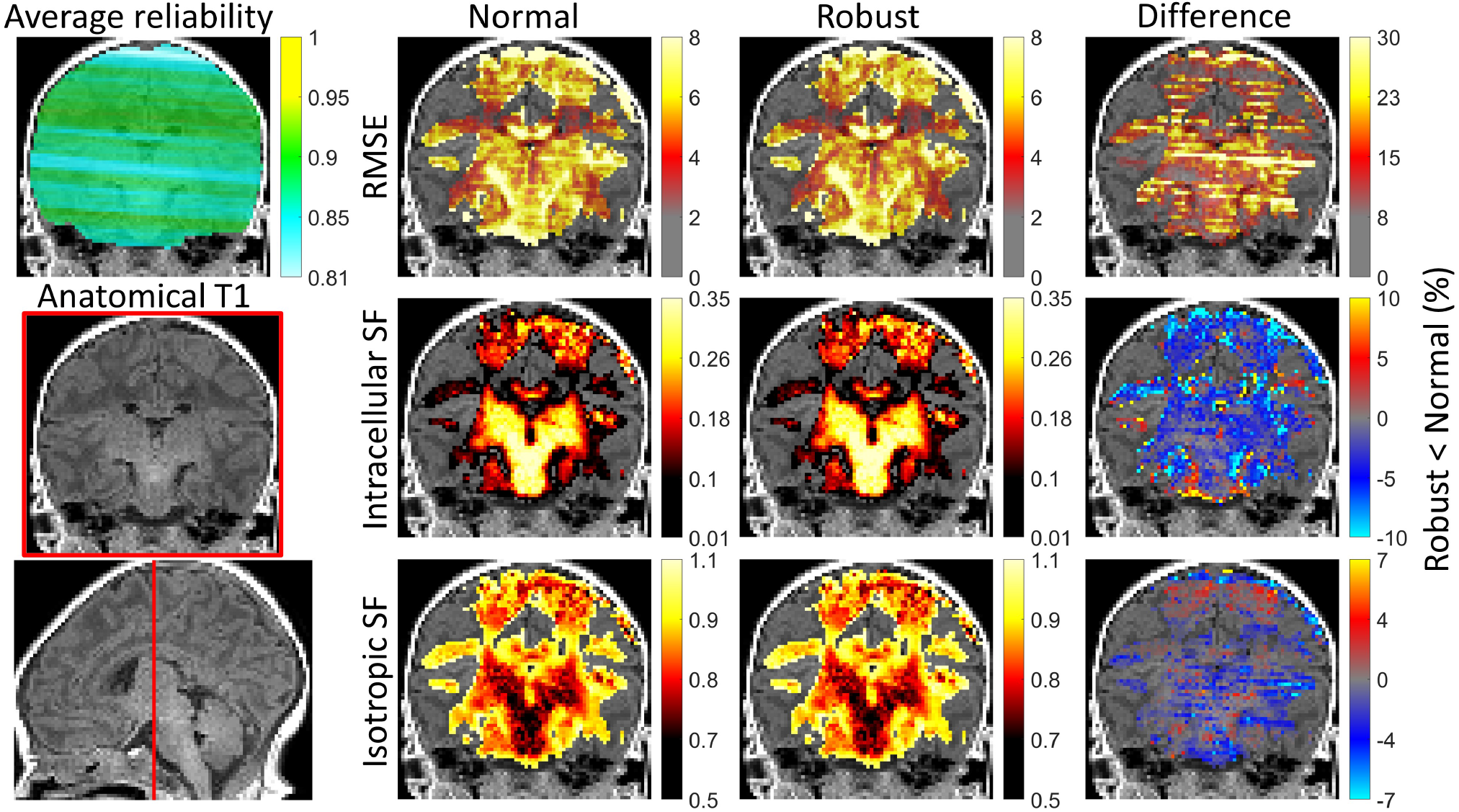
Summary of the evaluation with in vivo infant data. Coronal images visualize the slicewise artefacts that typically occur in the axial plane. Due to rotational subject motion (yaw, pitch, and roll), the acquired axial plane becomes an oblique plane during the image transformations that are necessary to align images to same spatial coordinates. On the left, an average reliability or confidence map and T1-image visualize the artefactual regions in the measurements after the image alignment. The bottom image in the left column shows a sagittal slice in which the red line indicates the position of the coronal slice. The three columns from the right are results for normal COMMIT, robust COMMIT_r and their difference, respectively. The first row details the results for root mean squared error (RMSE), the second row for intracellular signal fractions, and the third row for isotropic signal fractions. In this case, the signal drop outliers are seen as increased diffusivity in random directions and normal COMMIT tries to adjust for it by increasing isotropic signal fraction in the affected slices. This results in an inceased RMSE in the corresponding slices as well as slightly overestimated isotropic signal fraction which can is easiest to see in the corresponding difference map. This leads to interesting problem elsewhere in the brain (not affected by outliers) where normal COMMIT overestimates the intracellular signal fraction. It should be noted that this is a case example which likely cannot be generalized as the effect of outliers is difficult to predict and and depends on the affected gradient directions as well as the underlying brain structures.

The RMSE map of normal COMMIT was clearly affected by the outliers resulting in visible stripes in the image. On the contrary, the COMMIT_r RMSE map that describes the robust cost function does not have such stripes therefore the fitting is not affected by outliers. The difference RMSE map visualises the stripy pattern more prominently and ranges from 0 to 30%. The outlier effect on intracellular and isotropic signal fractions was less prominent in visual analysis i.e. less or no stripes. However, the difference between normal COMMIT and robust COMMIT_r depicts that the differences ranged from -10% to +10% even in regions that were less affected by outliers for intracellular signal fraction. For isotropic signal fraction the differences ranged from -7% to +7%.

## 4. Discussion

We demonstrated that tractogram filtering is severely affected by subject motion artefacts and that with our proposed robust augmentation these effects can be mitigated. In clinical research with uncooperative patients such as infants, it is highly likely that motion to some degree occurs during scanning. This leads to corrupted measurements which should not affect any modeling methods applied to the data. To best of our knowledge, this is the first time that motion related outliers are considered in the context of tractogram filtering therefore this update is crucial to enable tractogram filtering in clinical research.

The reason why we evaluated the proposed augmented cost function with simulated brains instead of real brain data was simply to ensure that nothing else in the relatively long dMRI processing pipeline might affect the results. For example, it is currently unknown issue, how outliers affect constrained spherical deconvolution based probabilistic tractographies. While there have been proposals for robust higher order model estimators (Pannek et al., 2012), such are not widely available. Furthermore, developing and evaluation of robustness of currently available constrained spherical deconvolution tractography algorithms are beyond the main scope of this study.

### Comparison to other filtering methods

While similar weighted cost function as in eq. 1 has been proposed before in SIFT filtering algorithm (Smith et al., 2013), those have been designed and tested to account for partial voluming related artefacts - not subject motion. The main difference in these artefact types is that partial voluming affects all dMRI data whereas subject motion affects only part of the dMRI data randomly. Therefore, adjusting for partial voluming requires one three-dimensional reliability image whereas adjusting for subject motion requires four-dimensional reliability image as the measurement reliability must be accounted for each dMRI data separately. This difference in the implementations of the algorithms also makes the accurate comparison of them fall outside the scope of this study.

### Correcting for artefacts

Our proposed algorithm (Fig. 1) can also be used to adjust for partial voluming but the necessity of that depends on the forward model used in COMMIT. For example, with ball and sticks model, voxels containing cerebrospinal fluid or gray matter can be described with an increased contribution from a ball compartment therefore the contribution of a stick compartment could be correct even without additional reliability weighting. If reliability weights are used, then the estimate for ball compartment would likely be improved but that should not still affect the filtered tractogram.

With motion induced artefacts, the outliers cause anisotropic signal deviations (Sairanen et al., 2017) affecting only part of the dMRI data. Therefore, COMMIT cannot adjust for those deviations simply by increasing the contribution of the ball compartment as the deviations are not isotropic over dMRI measurements. This is demonstrated in Fig. 10 where normal COMMIT obtains incorrect estimates for isotropic signal fraction maps i.e. ball compartments. Issue propagates causing also incorrect estimates for intracellular signal fraction maps i.e. stick contributions. Therefore, a local motion artefact can have a global adverse effect in tractography filtering if not accounted for.

### Statistical analysis

The global connectivity differences (Fig. 5) showed that normal COMMIT results varied heavily depending on the used outlier scheme (uniform or cluster) as well as on the outlier percentage (5% and 10%) whereas the robust COMMIT_r results remained relatively intact in all cases. Statistical tests 1 depicted that global connectivity was significantly affected by outliers with normal COMMIT producing also large effect sizes when compared against Baseline. On the contrary, comparison between Baseline and robust COMMIT_r resulted in small effect sizes despite two-sample Kolmogorov-Smirnov tests reporting statistically significant differences in 10% outlier simulations. It is possible that the amount of simulated outliers (%10) was already reaching the limit after which the missing data problem becomes too severe even for robust modeling methods. This could also be related to sample size being so large that Kolmogorov-Smirnov test finds any differences statistically significant despite having relatively small effect sizes. Therefore, in future studies some other non-parametric test could provide better results.

A more in-depth analysis of the connection from medulla to the right precentral gyrus (Figs. 6 and 7) revealed that ANOVA failed to find statistically significant differences between the groups with a p-value of 0.13 in 10% outlier simulations whereas significant differences were found in 5% outlier simulations with p-value less than 0.05. The non-parametric Friedman’s tests indicated for both outlier percentages that differences existed between the groups with a p-value less than 0.05. Notably, the effect sizes in comparison between Baseline and robust COMMIT_r remained much smaller than in comparisons between Baseline and normal COMMIT providing support for our proposed method being capable to mitigate these artefacts even for individual network edges.

In summary, it remains unsolved what test statistic would be the most suitable to analyse such data that is affected by outliers in anisotropic manner. We used two alternative approaches to evaluate the differences in group averages (ANOVA) and group distributions (Kolmogorov-Smirnov). Average based analyses are likely inefficient to locate all differences arising from outliers in the data whereas non-parameteric test can be even too sensitive to baseline shifts. Therefore, instead of statistical significance, the obtained effect sizes are likely more meaningful results.

### Robust modeling vs outlier replacement

This section extends outside the main scope of this study and is intended for the readers interested in slicewise outliers and how they should be addressed in dMRI in general. We added this section because we feel that the use of out-lier replacement in diffusion weighted literature is not truly justified and should not be continued in its current state. To understand our reasoning, readers are encouraged to familiarize the concept of outlier replacement which in statistics is known as data imputation. For this, we recommend the textbook *Statistical analysis with missing data* by Little and Rubin (Little, 2002).

Outlier replacement in dMRI is a form of multiple imputation which has been developed to correct for missing data in statistical analyses. Benefits of well performed imputation include decreased bias, increased precision, and most conveniently the ability to apply standard statistical tests and model estimators. For example, applying the standard t-test on sample that contains many missing or incorrect measurements could result in highly incorrect outcome whereas using a fixed sample that contains correctly imputed data could provide more reasonable results. This is, of course, the reason why outlier replacement seems so tempting in the context of dMRI: simply replace outliers and use the rest of the analysis pipeline as it is.

Imputation methods range from naive neighborhood interpolation (outlier is replaced e.g. by the average of its neighbors) and sample statistic based replacements (outlier is replaced by e.g. the mean or median value of the sample) to complex model prediction based replacements. Some of these ideas have already been transferred to dMRI usage by replacing outliers by their q-space neighborhood (Niethammer et al., 2007) or model based estimations (Lauzon et al., 2013; Andersson et al., 2016; Koch et al., 2019). These methods have in common that they depend on perfect outlier detection which can be problematic if there are many outliers. If some of the outliers are not correctly detected, this can lead into bias in the interpolation or modeling used in imputation which would propagate to the diffusion modeling that is performed using a normal estimator.

The idea in robust modeling is to account for the unreliability of the measurements and weigh each data point accordingly. In dMRI, such reliability can be obtained from voxelwise residuals (outliers tend to have large residuals) which has been implemented in algorithms suchs as RESTORE (Chang et al., 2005) and REKINDLE (Tax et al., 2015) for tensor model fitting. Similar approach can be applied to nearly any model.

Voxelwise outlier detection, however, is suboptimal in dMRI because artefacts in the echo-planar imaging result in whole slices being incorrect. Therefore, detecting subject motion related outliers in slicewise manner and assigning the reliability to all voxels in those slices is arguably be more powerful approach (Andersson et al., 2016; Sairanen et al., 2018). Moreover, slicewise outliers can be used as complementary information for voxelwise estimators to adjust for more local sources of uncertainties e.g. pulsation due to heartbeat.

To illustrate the performance of the aforementioned ideas, we provide a minimal example in which we compare naive outlier replacement to simple robust model estimation. We used the constrained spherical deconvolution (CSD) algorithm implemented in DIPY (Tournier et al., 2007; Garyfallidis et al., 2014) in this evaluation as evaluating the differences using COMMIT framework would be computationally inefficient (and well beyond the purpose of the current study). We reason that if this simpler analysis cannot provide support for using outlier replacement, there is little reason to test it in a more complex analysis.

In CSD, we used the default DIPY-library parameters with *L*_*max*_ 8, tau 0.1, 362 vertices on the symmetrical sphere, relative peak threshold of 0.5, minimal peak difference angle of 25 degrees, and 50 iterations. We developed four streamline setups that are described in Fig. 11. All setups consisted of three axial slices in which the middle slice was affected by a full signal dropout artefact. We investigated what happens to CSD signal prediction if *i*) nothing was done to the outlier, *ii*) outlier was replaced with a naive neighborhood interpolation, and *iii*) spherical harmonic coefficients used in CSD were obtained using a robust in-house version of CSD algorithm. The in-house algorithm simply decreased the outlier weight to zero in the linear least squares estimation of the spherical harmonic coefficients.

**Figure 11:**
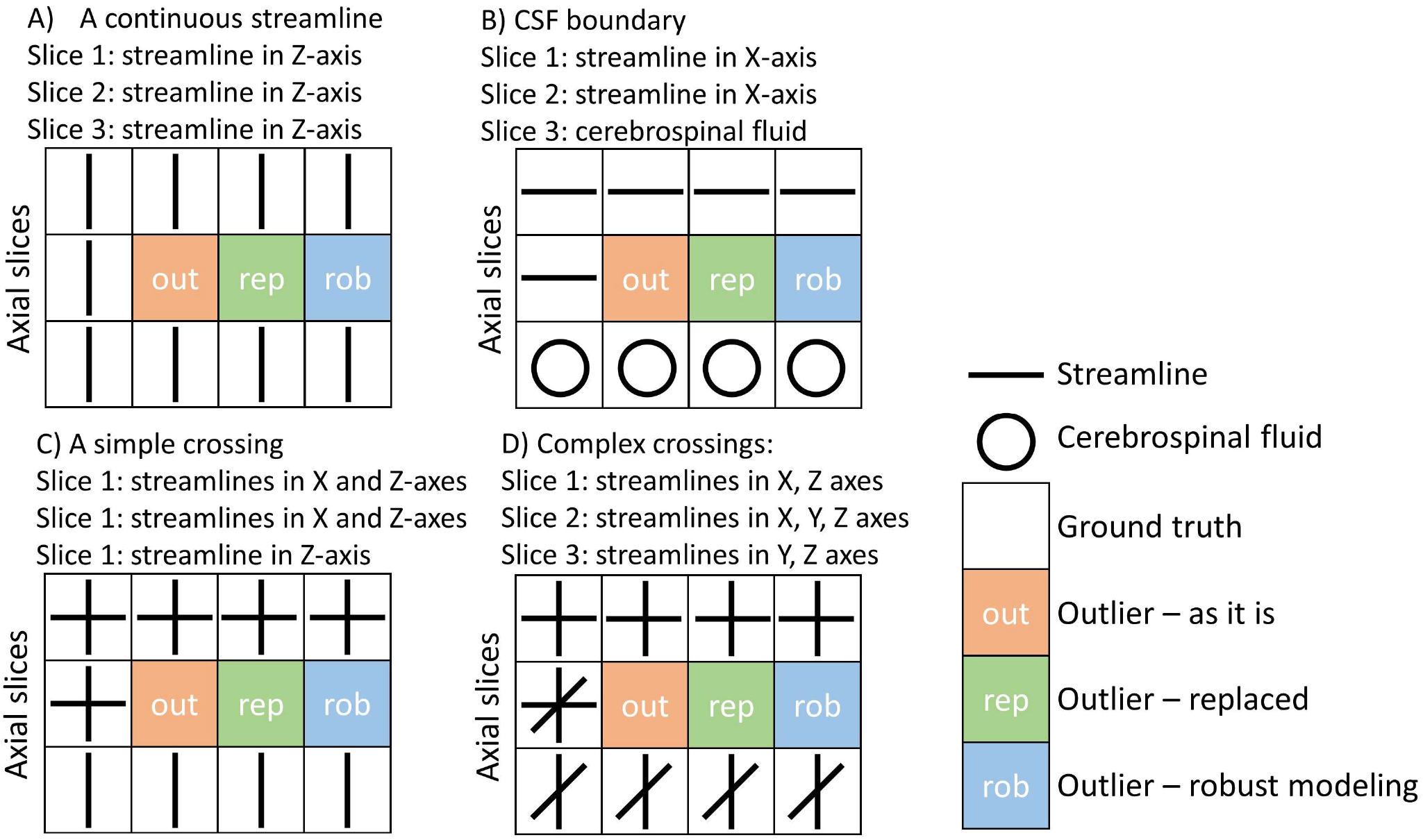
Illustration of the streamline phantom setups used to investigate differences between outlier replacement and robust modeling. The left column in all setups is a ground truth with no outlier and normal modeling, the second column is a control with outlier and normal modeling, the third column is replaced outlier with normal modeling, and the fourth column is outlier and robust modeling. Outlier replacement was calculated as an average from the slice below and above the outlier (i.e. spatial neighborhood). A) A single streamline traverses in parallel to z-axis throughout the phantom. Therefore, the outlier replacement produces exactly the missing data and should provide the same result with the ground truth. B) A single streamline traverses parallel to x-axis in two upper slices with the third slice consisting of cerebrospinal fluid. C) Two upper slices consist of streamline crossings in x and z-axes with the third consisting of streamline traversing only parallel to z-axis. D) The first slice consists of streamline crossings in x and z-axes, the middle slice consists of crossings in all main axes, and the third slice in crossing in y and z-axes.

We used infinite signal-to-noise ratio to evaluate only the effects of the signal dropout. Outliers were introduced incrementally from 0 to 9 of one of the HCP gradient scheme shell with b-value of 2000*s*/*mm*^2^ using three different schemes in outlier selection. The first scheme represented the worst possible situation where outliers were clustered in the *q*-space (e.g. Fig. 4), the second scheme represented the best possible situation where outliers were uniformly placed in the *q*-space based on their electrostatic repulsion (Sairanen et al., 2017), and the third scheme represented randomly selected outliers. In the first two cases we evaluated all possible 90 cases whereas the random scheme consisted of 500 combinations. Signal predictions from these schemes were compared to prediction from the ground truth fit that was not affected by outliers. In this simulation we did not add Rician noise, therefore direct comparison to ground truth is sound.

It should be noted that random outlier replacement is a very poor method to evaluate these effects in dMRI due to the extremely large number of possible combinations and we performed it only for illustrative purposes. For example, selecting 9 outliers out of 90 diffusion weighted images can be performed in 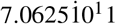 different ways therefore selecting 500 of these randomly might not represent the population well. Since evaluating all possible combinations is possible only for a smaller number of gradient directions (Sairanen et al., 2017), it is far more informative to investigate the best (uniform) and the worst (clusters) extreme cases.

Results of these four setups, shown in Fig. 12 demonstrate that if nothing is done to the outliers, the difference to ground truth increases linearly as the number of outliers increase. Of course, after the mathematical problem becomes ill-conditioned a more chaotic results would be expected (Sairanen et al., 2017) but this is likely to occur with much larger number of outliers. Based on the Fig. 12 the naive outlier replacement outperformed robust modeling only in the Setup A in which the outlier was replaced by identical information from the neighboring voxels. In setups B, C, and D with more complex and perhaps realistic streamline combinations robust modeling outperformed outlier replacement by providing results that were closer to the ground truth.

**Figure 12:**
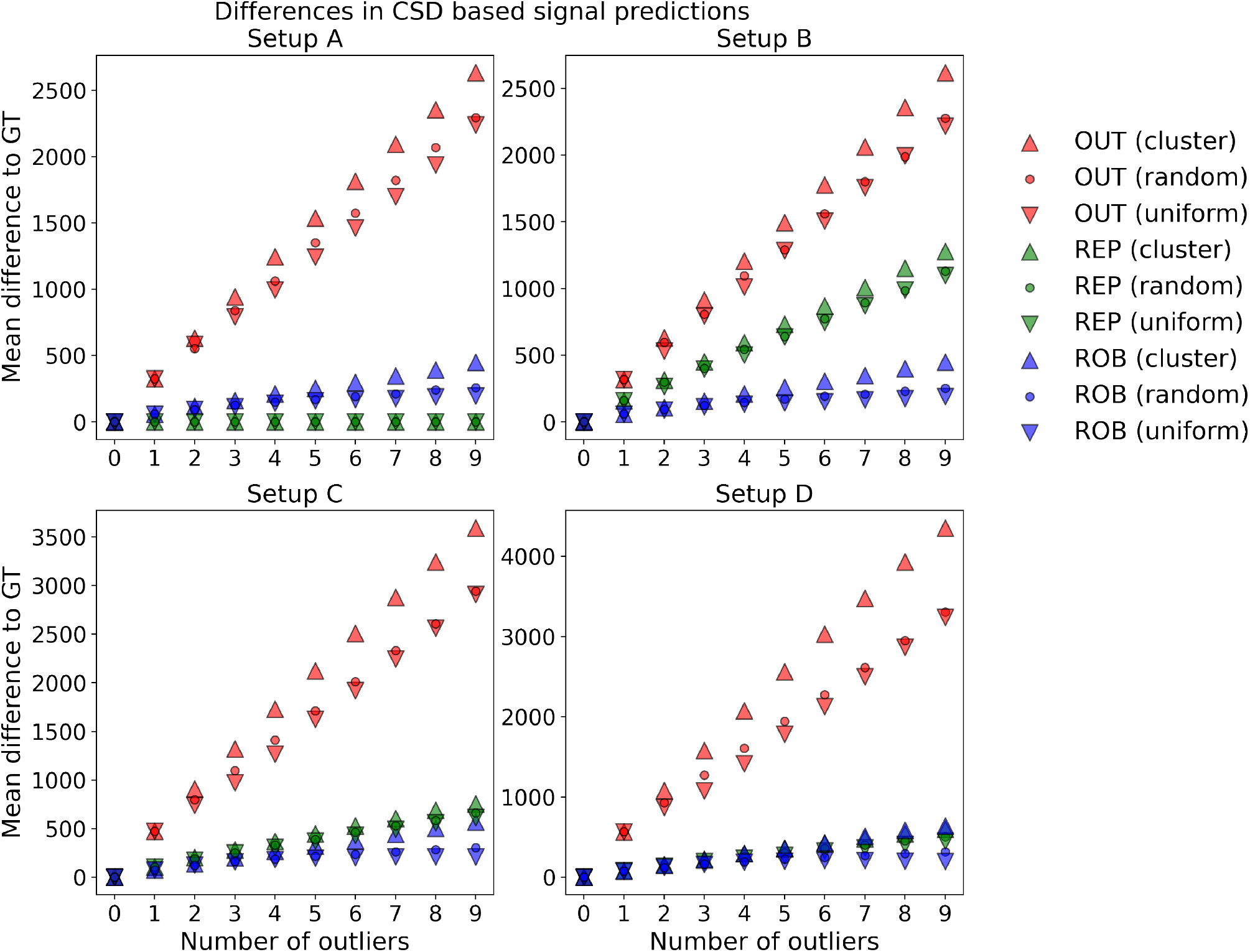
Outlier replacement compared to robust modeling in constrained spherical deconvolution (CSD). Upward triangles (△) indicate clustered outliers, dots (∘) indicate randomly placed outliers, and downward triangles (▽) indicate uniformly placed outliers. All values are calculated as an average residual from ground truth (GT) signal prediction. Red markers (OUT) are results from normal CSD, green markers (REP) are results from normal CSD with outlier replacement, and blue markers (ROB) are results from the robust CSD. The clustered outliers (△) result in the largest differences in all cases and the uniform outliers (▽) result in the smallest differences. Random cases (∘) are generally closer to the uniform situation as the extreme outliers tend to result in heavily tailed distributions. Outlier replacement provided better results only in the Setup A in which the outlier could be replaced with similar data that was missing. In all other cases, where the spatial neighborhood did not exactly represent the missing measurement, the outlier replacement produced larger difference to the ground truth than the robust estimator. As the complexity of the streamline phantom increases in Setups C and D, the difference between replacement and robust methods become smaller. Importantly, both the robust modeling and outlier replacement improved the CSD prediction compared to the baseline case (OUT).

It can be reasonably argued that the rather naive outlier replacement we implemented here could be improved with already available proposals of q-space neighborhood (Niethammer et al., 2007) or model based estimations (Lauzon et al., 2013; Andersson et al., 2016; Koch et al., 2019). However, same applies to the in-house robust spherical harmonic linear least squares estimator we developed for this task which simply down weighs the outlier measurements. However, while it might be possible to fine tune outlier replacement in dMRI to the degree that matches the robust modeling, outlier replacement would still lack the ability to evaluate the uncertainty in the fitted model rendering it less useful method for clinical usage that might require or benefit from knowledge of the method’s uncertainty (e.g. surgery or radiotherapy).

The reader might be confused by the previous statement that imputation could not contain information about uncertainties while even the text book by Little and Rubin (Little, 2002) we cited has a chapter called “*Estimation of Imputation Uncertainty*”. To understand this, remember that the imputed sample in dMRI is generally an axial slice in a three dimensional stack of slices that are a part of four dimensional series of diffusion weighted images. MRI scan of the whole series takes several minutes during which the patient’s head tends to move and especially rotate (yaw, pitch, and roll). These rotated images must be aligned with some reference image before model fitting but by doing so the image registration transforms the axial (outlier) slice into an oblique plane which is an interpolation between the slice and its neighbors.

This process of image alignment is described in Figure 1. of Sairanen et al. 2018 (Sairanen et al., 2018) but in short, afterwards it is likely impossible to accurately distinguish an imputed signal fraction from a normal signal. This means that likelihood based estimators or bootstrap methods (Whitcher et al., 2008) no longer can estimate the uncertainty of the fitted diffusion model. On the contrary, robust modeling that is based on measurement reliability weights would still be able to tell this difference. Therefore, any clinical application that might benefit of these uncertainty estimates would be hindered by using outlier replacement. To avoid such bottleneck in the future of dMRI, we argue that it would be highly beneficial for the dMRI community to avoid using the outlier replacement in its current form. Even in basic neuroscience, it could be beneficial to know the voxelwise distributions of model derived values such as fractional anisotropy (e.g., *FA* = 0.6 ± 0.03) to perform sound statistical analyses.

### Where to go from here?

We considered only post-scan motion corrections in this study because during-scan corrections should be able to produce data that does not need these correction algorithms. The problem with during-scan corrections is their limited availability due to external hardware requirements or still experimental software. Due to the long time span of tens of years required to advance MRI technology in clinical use, it is unlikely that these during-scan correction methods would be so widely available in clinical research centers that postscan corrections such as our proposal are rendered obsolete any time soon. While the post-scan corrections are more like a remedy to the symptom instead of cure to the cause, novel studies on clinical patients and even infants are increasingly proposed and carried out therefore the need for robust tools is current and cannot wait decades for hardware based solutions.

## 5. Conclusion

We proposed a augmentation to a tractogram filtering algorithm COMMIT that renders it robust towards subject motion caused outliers in the measurements. This addition is necessary for conducting tractogram filtering in clinical research where subject motion is often unavoidable. While robust data processing has been implemented before in the context of diffusion tensor and higher order model estimations, it has not been previously implemented for tractogram filtering. We used realistic whole brain Monte-Carlo simulations that account for kissing and crossing fiber structure as well as partial volumeing to successfully demonstrate that our augmentation is capable to accurately map the structural brain connectivity in the presence of such outliers in the data. We also demonstrated that if this correction is not done, the structural connectivity estimates can become strongly biased. With this update any clinical study investigating structural connectomics of children or uncooperative patient populations can robustly perform their analyses without the need to exclude subjects with outliers from them.

## Acknowledgements

V.S. was supported by the Brain Research Foundation Verona, Orion Research Foundation sr, and Instrumentarium Science Foundation sr. M.O-P., S.S. and A.D have no relevant financial or non-financial interests to disclose. C.G. was funded by the Swiss National Science Foundation (SNSF) grant PP00P3_176984, the Stiftung zur Förderung der gastroenterologischen und allgemeinen klinischen Forschung, EUROSTAR E!113682 HORIZON2020. The authors wish to thank the Finnish Grid and Cloud Infrastructure (FGCI) for supporting this project with computational and data storage resources.

## Footnotes

1 https://github.com/daducci/COMMIT

